# In situ evidence for systematic membrane thickness variation across cellular organelles

**DOI:** 10.1101/2025.03.13.642912

**Authors:** Desislava Glushkova, Stefanie Böhm, Martin Beck

## Abstract

In eukaryotes, membrane-bound organelles create distinct molecular environments. The compartmentalizing lipid bilayer is a dynamic composite material, whose thickness and curvature modulate the structure and function of membrane proteins. In vitro, bilayer thickness correlates with lipid composition. Cellular membranes in situ, however, are continuously remodeled and the spatial variation of their biophysical properties remains understudied. Here, we present a computational approach to measure local membrane thickness in cryo-electron tomograms. Analysis of *Chlamydomonas reihardtii* and human cells reveals systematic thickness variations within and across organelles. These findings orthogonally support models of hydrophobic matching for differential sorting of proteins based on their transmembrane domain lengths, e.g., across the Golgi apparatus. Our workflow is computationally efficient, public, easy to integrate within existing tomogram analysis pipelines, and enables membrane thickness measurements across experimental conditions. Using this approach, relationships between membrane composition, thickness, and function are explored in situ, with broader applications across membrane biology.

## Introduction

Cellular membranes are composite materials, consisting of a lipid bilayer interspersed with membrane proteins, which serve as essential sites for biochemical processes in all living organisms [1]. The lipid composition of membranes affects their biophysical properties, including curvature [2], thickness [3][4], fluidity [4][5], and compressibility [2][4]. Both direct lipid-protein interactions and indirect changes in membrane properties regulate the function of membrane proteins and ultimately affect diverse biological pathways [6][7][8][9][10][11][12].

Eukaryotic cells invest substantial resources in generating a diverse lipidome, comprising more than a thousand distinct species [13], with multiple lipid metabolic pathways underlying the diversity of membrane lipid composition across species [14][15], tissues, and cell types [15][16][17]. Lipid composition can also change dynamically during various cellular processes. For example, HeLa cells actively regulate both the composition and the spatial distribution of lipids during cell division, which alters the mechanical properties of their membranes compared to non-dividing cells [18]. Similarly, hormonal tissue differentiation involves modifications to the lipidome of epithelial cells, including changes in the phospholipid profiles [19] and shifts from sphingomyelin to glycosphingolipid [20].

Beyond cell-type differences, organelle membranes within individual cells possess distinctive lipid “fingerprints”, likely related to their specific functions. Lipidomic analyses of subcellular fractions have revealed non-homogeneous distributions of phospholipids and sterols across eukaryotic cellular compartments [21][22], with some lipids being primarily found in specific organelles, e.g., cardiolipin in the inner mitochondrial membrane [23] and lysobisphosphatidic acid in late endosomes [24]. Although the endoplasmic reticulum (ER) serves as the primary site for phospholipid and sterol synthesis, its own membranes contain relatively low concentrations of the latter, as cholesterol is rapidly shuttled through the secretory pathway to the Golgi, where sphingolipids are synthesized. Both sphingolipids and cholesterol then get transported to the plasma membrane (PM), where they contribute to creating a tightly-packed, relatively impermeable barrier around the cell [13][21]. Endosomal membranes exhibit a compositional gradient during maturation: early endosomes are thought to resemble the PM in their lipid profile, while sterol and phosphatidylserine levels progressively decrease during progression into late endosomes [22].

Despite significant advances in understanding lipid distribution across cell types and cellular compartments, lipidomic approaches face several limitations: cell lysis eliminates spatial information on membrane organization; obtaining pure organelle fractions can be challenging, particularly for membranes of similar size and density; and the analytical outcomes are sensitive to variations in lipid extraction protocols [21]. Furthermore, the more elaborate morphology of many organelles, exemplified by the cisternae of the Golgi apparatus, is lost during isolation.

Organelle-specific lipid compositions not only define membrane identity, but also create distinct physicochemical environments that influence protein function and intracellular trafficking. The Golgi apparatus serves as a compelling example: it has been proposed that variations in membrane thickness drive the partitioning of transmembrane proteins into distinct Golgi sub-compartments [25][26], thus minimizing hydrophobic mismatch and its associated energetic penalty [27][28]. According to this model, proteins with longer transmembrane spans preferentially partition to the thicker cholesterol-rich trans Golgi region, from which they are subsequently directed to the PM, while proteins with shorter transmembrane spans are retained in the earlier, thinner Golgi membranes [29][30].

Multiple experimental approaches have established a clear relationship between membrane thickness and lipid composition. Small-angle neutron scattering (SANS) studies have characterized the thicknesses of both the hydrophobic cores and hydrated layers of vesicles with various lipid compositions, demonstrating that hydrophobic core thickness increases with acyl chain length, while the water layer thickness remains relatively constant at approximately 1.5-1.8 nm [31][32]. Small-angle X-ray scattering (SAXS) analyses have similarly demonstrated a linear relationship between the thickness of the hydrophobic core and the acyl chain length in pure phosphatidylcholine vesicles [33].

More recently, cryo-electron microscopy (cryo-EM) has enabled direct visualization of membrane thickness variations at high resolution [34][35][36]. The bilayer thicknesses of vesicles with defined lipid compositions have been measured from trough-to-trough distances in 2D-projected intensity profiles from both micrographs [34][36] and tomograms [35], confirming the linear relationship between acyl chain length and membrane thickness [34]. In phase-separating lipid mixtures, segregation of membrane microdomains into liquid-ordered and liquid-disordered regions has been implied based on fitting the thickness distributions of individual vesicles with bimodal Gaussian models [34][35]. The bilayer thicknesses reported in the cryo-EM studies align well with the dimensions of the hydrophobic core previously established by SANS and SAXS measurements [31][32][33]. Although cryo-EM provides direct visualization of membranes at high-resolution, it has primarily been applied to measure the thicknesses of synthetic bilayers with reduced compositional complexity compared to native cellular membranes. Furthermore, existing approaches typically employ 2D analysis methods that do not account for variations in membrane curvature and, when relying on manual tracing and measurement, have limited throughput.

The converging evidence for organelle-specific lipid distributions and the well-documented relationship between lipid composition and membrane thickness suggest that cellular membranes may exhibit characteristic thickness profiles, related to their functions. However, membrane thickness variations within cellular environments have largely remained uncharacterized. To this aim, we developed a semi-automated computational workflow to systematically measure bilayer thickness from cryo-electron tomogram segmentations of diverse cellular membranes. Our approach performs 3D thickness measurements in a voxel-wise manner, thus accounting for membrane curvature. The resulting membrane thickness maps can be overlaid with the original tomogram or with other structural features to provide biological context. By applying this workflow to a large publicly available *Chlamydomonas reinhardtii* dataset [37], we identified consistent thickness differences not only between organelle membranes, but also within the bounds of a single organelle. Measurements in a human cell line yielded similar results and showed that membrane thickness is affected by acute changes in lipid composition.

## Results

### Outline of the computational workflow for in situ membrane thickness analysis

Membrane thickness is a factor that directly influences the organization, structure, and function of membrane-associated proteins. Methods to reliably assess this parameter within a cellular context are therefore critical to understanding protein function at a molecular level. To date, cryo-electron tomography (cryo-ET) provides the highest-resolution direct visualization of native cellular structures within three-dimensional volumes [38]. Membrane structures within cells exhibit variable morphologies and potentially heterogeneous thicknesses that may reflect differences in their biophysical or functional properties. To explore this question, we developed a computational workflow for semi-automated measurement of bilayer thicknesses from membrane segmentations of cryo-electron tomogram data (Figure 1). Our approach allows for quantitative assessment of thickness variations both within individual membranes and across organelles (or species), as well as for correlation of membrane thickness with the coordinates of protein complexes contained in the same tomogram.

**Figure 1.**
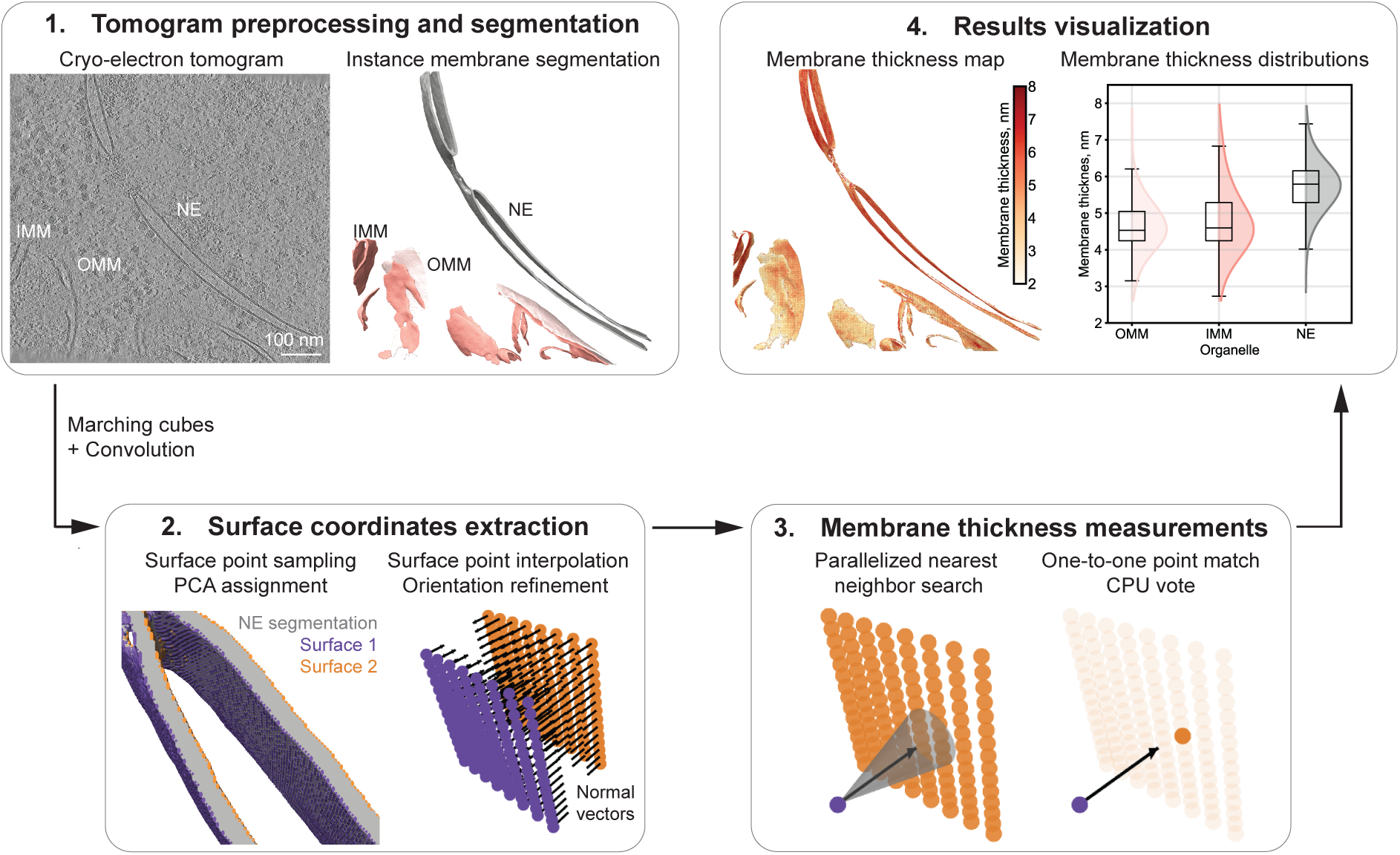
Computational workflow for membrane thickness analysis from cryo-electron tomograms. **1. Tomogram preprocessing and segmentation:** Reconstructed cryo-electron tomograms are processed with MemBrain-seg [39] to generate instance segmentations of different membranes in a given volume (here, NE: nuclear envelope, OMM: outer mitochondrial membrane, IMM: inner mitochondrial membrane). **2. Surface coordinates extraction:** A convolution-constrained marching cubes algorithm [43] outputs the coordinates and orientations of surface points. The segmentation surfaces are separated via principal component analysis (PCA, visualized with purple and orange points) and the normal vectors (black arrows) are refined through local weighted averaging. Additional surface-constrained point interpolation can increase the coverage. **3. Membrane thickness measurements:** For each point on one surface, a ray is projected along its normal vector. Parallelized cone search identifies potential nearest neighbor candidates on the opposite surface. The final one-to-one point match is determined through CPU-based voting. Membrane thickness is calculated as the Euclidean distance between matched point coordinates. **4. Results visualization:** The output files include membrane thickness maps (color intensity represents local membrane thickness), which can be used for co-localization with the coordinates of membrane-bound proteins, as well as data on thickness distributions across different membrane types.

The computational workflow is visualized in Figure 1. It begins with instance segmentation of individual membrane structures in the volume using MemBrain-seg [39] (Figure 1-1). Prior to segmentation, the tomograms can be denoised (e.g., with cryoCARE [40]) or filtered with a Wiener-like deconvolution filter (such as implemented in Warp [41]) to improve contrast and segmentation accuracy. Post segmentation, the output labels can be manually curated with visualization tools such as napari [42]. Unlike semantic membrane segmentation, where each voxel is classified as membrane or non-membrane (binary classification), in instance segmentation, voxels containing different membrane structures are assigned unique integer values. We then use the instance segmentation volume and the generated labels as inputs to perform targeted thickness measurements on specific membranes of interest within the tomogram.

In the second step, the coordinates and orientations of surface points are extracted from the segmentation volume (Figure 1-2). Initial surface points are generated using the scikit-image implementation of the marching cubes algorithm [43], coupled with a 3D convolution kernel that identifies edge voxels. For comprehensive coverage, additional points can be interpolated between existing ones, with the constraint that all points must lie on the actual segmentation boundary, as determined by the convolution kernel. The marching cubes algorithm assigns orientations to each surface point, which are then refined through weighted averaging within a specified neighborhood radius. Refining the orientations ensures smooth transitions across the segmentation surface, while still taking native membrane curvature into consideration. Sampled points are assigned to the “inner” or “outer” membrane surfaces (leaflets) through a computationally-efficient principal component analysis (PCA) of the global points orientations.

**Figure 2.**
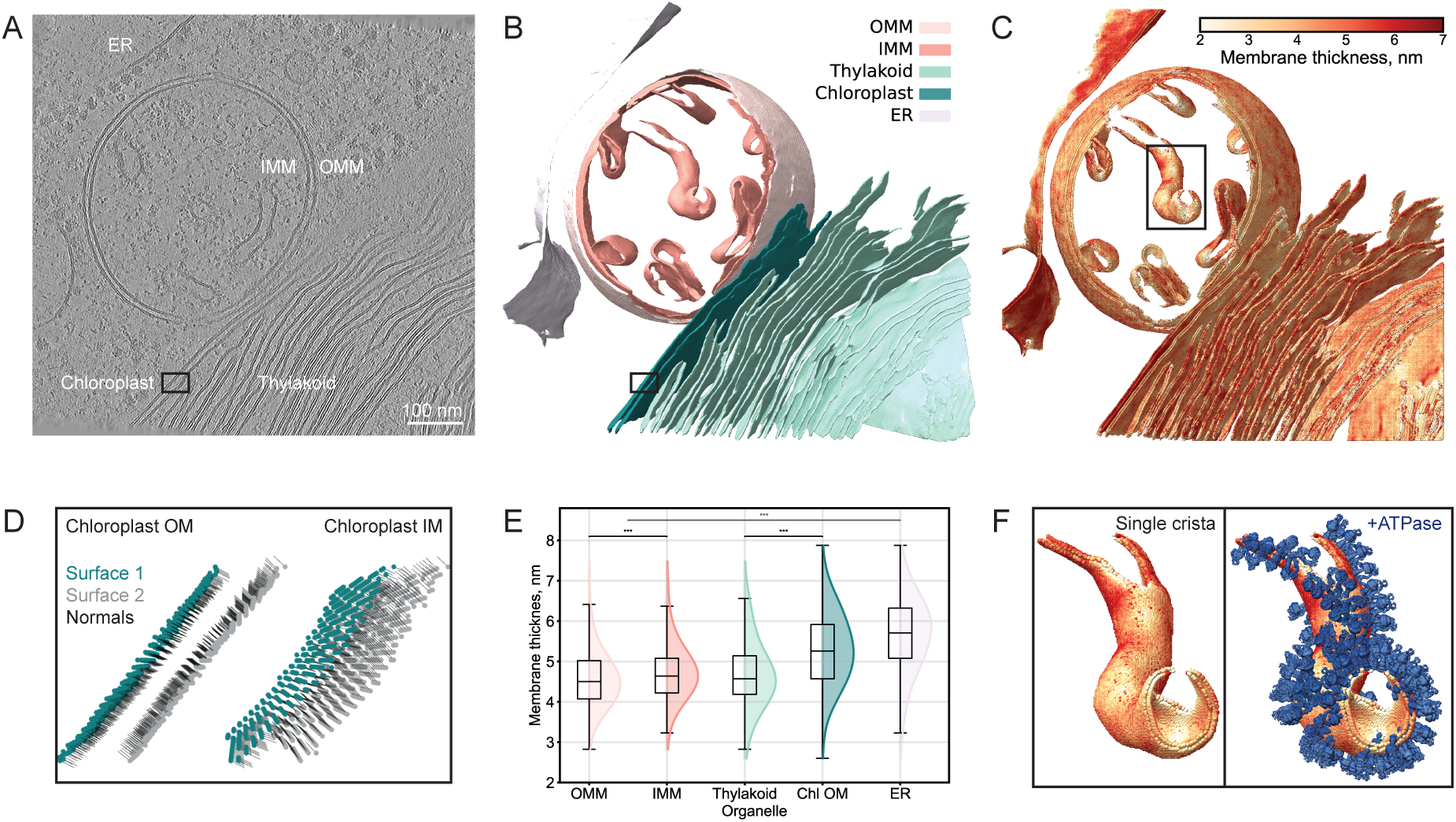
Analysis of organelle membrane thickness heterogeneity within an exemplifying tomogram. **A** Central slice from a denoised *Chlamydomonas reinhardtii* tomogram [37] (EMPIAR-11830, 2140.mrc, 7.84 Å/pix at bin4), showing the endoplasmic reticulum (ER), inner and outer mitochondrial membranes (IMM and OMM), thylakoid and chloroplast membranes. Scale bar, 100 nm. **B** Instance segmentation generated by MemBrain-seg [39], with distinct colors marking different membrane instances. **C** Membrane thickness map, where color intensity represents differences in local membrane thickness. **D** Detailed view of the surface point extraction and normal vector orientation step, shown for a region of the chloroplast membranes (marked by black boxes in A,B). Points assigned to surface one are colored in dark green, while points assigned to surface two are shown in light gray. Black lines indicate the normal vectors pointing from each point towards the opposite segmentation surface. **E** Comparative thickness distribution plots across membranes (color-coded as in B). The Kolmogorov-Smirnov (KS) test was used to assesses differences between the following distributions: OMM vs IMM (black line and symbols, ***: p < 0.001), thylakoid vs chloroplast outer membrane (Chl OM, black line and symbols, ***: p < 0.001), ER vs both mitochondrial membranes (gray line and symbols, ***: p < 0.001). Additionally, the Wasserstein distances and effect sizes within 95% confidence intervals were calculated. Detailed information on the outcomes of the statistical analysis can be found in Supplementary Table 1. **F** Co-localization of local membrane thickness with macromolecular complexes, demonstrated for a single mitochondrial crista (marked by a black box in E). ATP synthase complexes (blue, right panel) appear to localize to regions of the membrane with high curvature and/or smaller thickness. ATP synthase particle positions were obtained from a publicly-available repository [44] as determined by subtomogram-averaging in [37], with the resulting subtomogram average map (EMD-52802) used for visualization. A more detailed visualization of thickness variations and ATP synthase localization is provided in Supplementary Video 1.

In the third step, membrane thickness is measured for each labeled instance in the volume (Figure 1-3). For each point on one segmentation surface, a ray is projected along its normal vector direction. A cone search, parallelized either across GPU nodes or multiple CPU threads, identifies potential nearest neighbor candidates on the opposite segmentation surface. A CPU-based vote determines the final one-to-one point pair in order to prevent multiple matches for a given point that can cause inaccuracies in the thickness measurements. Membrane thickness is calculated as the Euclidean distance between the coordinates of the paired points (in voxels, afterwards scaled by the segmentation pixel size to output the distance in nm), with the normal-guided search accounting for the local curvature in 3D space.

**Figure 3.**
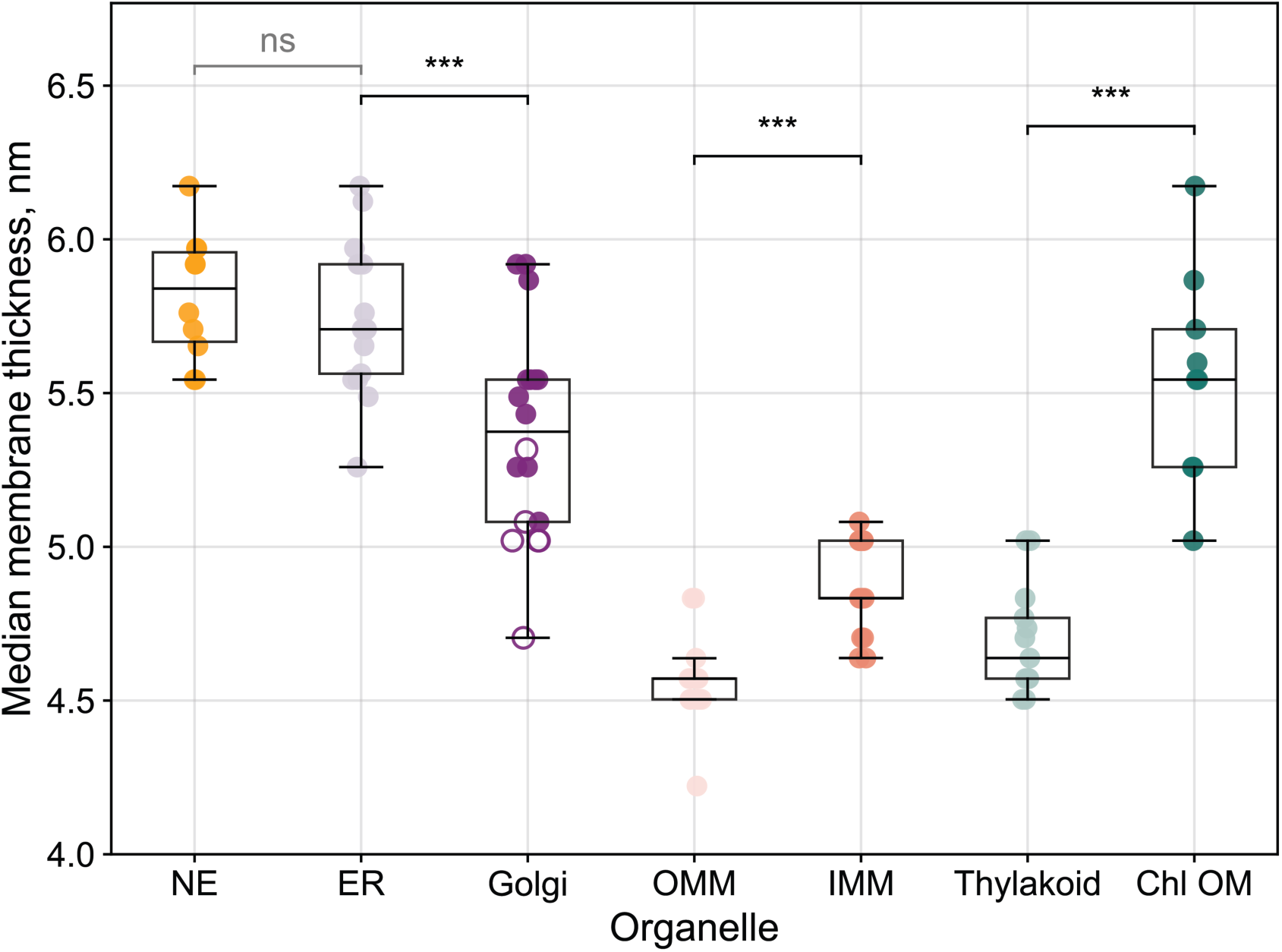
Organelle-specific thickness patterns across tomograms. Summary of median membrane thicknesses across 51 *Chlamydomonas reihardtii* tomograms [37] (EMPIAR-11830). Each point corresponds to the median thickness value for a given membrane within a single tomogram: nuclear envelope (NE, n=10, 2.1M measurements), endoplasmic reticulum (ER, n=18, 2.6M measurements), outer mitochondrial membrane (OMM, n=19, 4M measurements), inner mitochondrial membrane (IMM, n=19, 10M measurements), thylakoid (n=14, 26M measurements), chloroplast outer membrane (Chl OM, n=9, 1.2M measurements). For the Golgi apparatus (n=6 tomograms), three measurements per tomogram are displayed to capture the thickness gradient across cisternae (addressed in more detail on Figure 5): cis and trans Golgi network membranes (filled circles) and medial Golgi membranes (empty circles). Statistical comparisons of the median values were performed for functionally-related membranes: NE vs ER (unpaired t-test, ns: not significant, p = 0.439), ER vs medial Golgi (paired t-test, ***: p < 0.001), OMM vs IMM (paired t-test, ***: p<0.001), and thylakoid vs Chl OM (paired t-test, ***: p < 0.001).

Finally, the results are outputted in multiple visualization formats for both qualitative and quantitative analysis (Figure 1-4). The membrane thickness maps, where color intensity corresponds to local thickness, provide an intuitive visualization of local thickness variations across different membrane regions. When exported as particle motive lists, the thickness maps can be used for contextual analysis through co-localization with the coordinates of membrane-bound proteins or other structures of interest. For quantitative comparisons, the thickness distributions of each membrane instance can be plotted and analyzed using statistical tests to evaluate differences across individual membranes, organelles, or experimental conditions. The visualization options allow for straightforward comprehensive analysis of membrane thickness in different cellular contexts.

**Figure 4.**
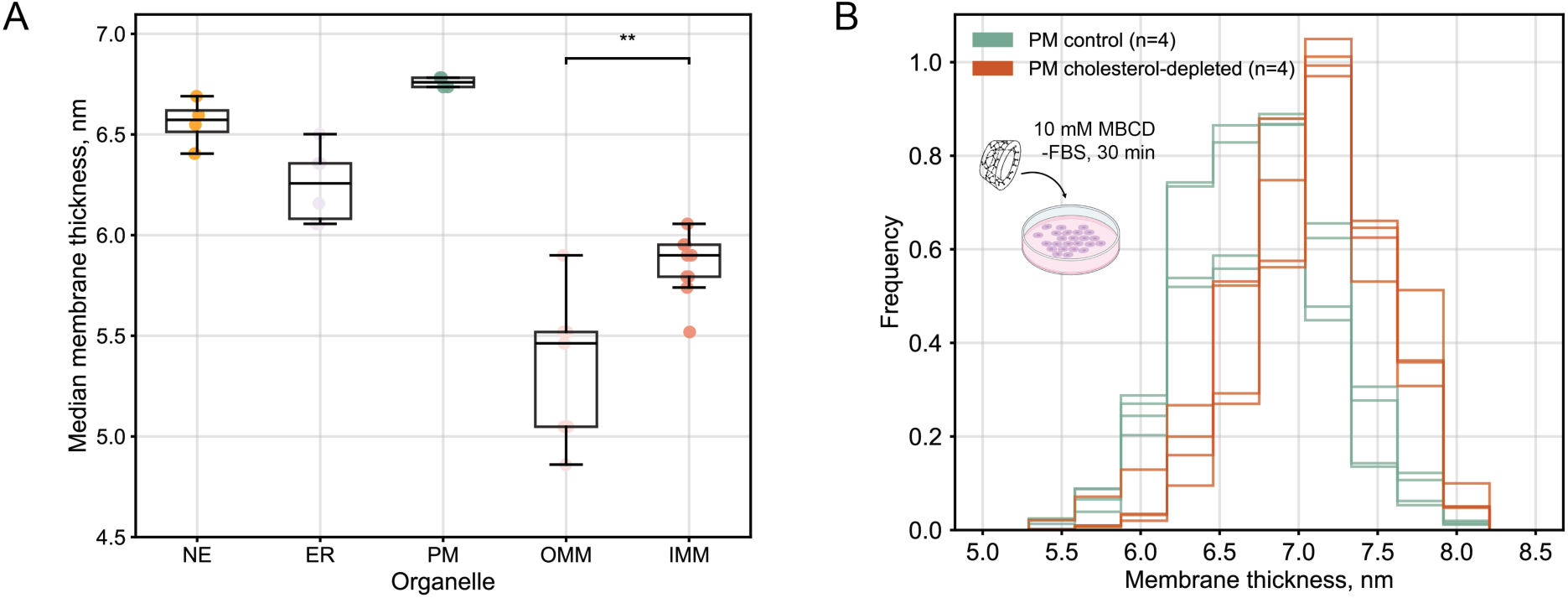
Organelle-specific thickness patterns are consistent in human cells. **A** Analysis of median thicknesses across membrane types in human embryonic kidney cells (HEK293): nuclear envelope (NE, n=4, 0.5M measurements), endoplasmic reticulum (ER, n=6, 0.4M measurements), plasma membrane (PM, n=4, 0.4M measurements), and outer and inner mitochondrial membranes (OMM, n=9, 0.9M measurements and IMM, n=9, 3.2M measurements). Statistical significance between the OMM and IMM median thicknesses was assessed by paired t-test per tomogram (**: p = 0.002). No unpaired t-tests for NE vs ER or NE/ER vs PM were performed due to the small number of membrane instances. **B** Validation of the thickness estimation sensitivity via a cholesterol depletion experiment. HEK293 cells were treated with methyl-beta-cyclodextrin (MBCD, 10 mM, 30 min) to extract plasma membrane cholesterol. Normalized thickness distributions of plasma membranes from control (n=4) and cholesterol-depleted (n=4) cells demonstrate that a change in lipid composition leads to a detectable difference in membrane thickness.

### Membrane thickness measurements applied to in situ experimental data

To test the performance of our workflow on in situ experimental data, we applied it to a publicly available denoised tomogram of the green algae *Chlamydomonas reinhardtii* [37] (Figure 2A).

Using MemBrain-seg’s connected component analysis [39], we generated an instance segmentation volume where distinct voxel values corresponded to different organelle membrane identities, including the rough endoplasmic reticulum (ER), inner and outer mitochondrial membranes (IMM and OMM), thylakoid, and chloroplast envelope membranes (Figure 2B). From this segmentation, we computed a membrane thickness map that captured local thickness variations (Figure 2C). To do so, we extracted the coordinates and orientations of surface points and classified them into opposing membrane surfaces, as demonstrated for a section of the chloroplast membranes, where the normal vectors point towards the opposite segmentation surface (Figure 2D).

Hundreds of thousands of individual distance measurements per membrane instance yielded highly-representative thickness distributions (Figure 2E). We observed statistically-significant thickness differences between organelle membranes within the same tomogram (with projection images acquired under identical imaging conditions). The IMM and OMM differed by 0.2 nm (Wasserstein distance), with highly-significant distribution differences (Kolmogorov-Smirnov test, p < 0.001) and a small but consistent effect size. We observed more pronounced differences between the thylakoid and chloroplast outer membranes (0.6 nm Wasserstein distance, p < 0.001, medium effect size). Notably, both mitochondrial membranes were significantly thinner than the ER (p < 0.001), with a large effect size. Complete statistical analysis with confidence intervals is provided in Supplementary Table 1.

Beyond quantitative comparisons, we used the membrane thickness maps for contextual analysis of the coordinates of ATP synthase complexes (determined through subtomogram averaging in [37]). By overlaying known particle positions, we visually assessed that the complexes appear to localize to regions of high curvature and/or decreased thickness within the cristae (Figure 2F and Supplementary Video 1 for more detailed visualization). This spatial correlation represents just one example of potentially interesting relationships between membrane thickness, curvature, and the structural organization of membrane protein complexes.

To assess the robustness of the computational workflow to different inputs, we compared the membrane thickness measurements on the same raw data with or without applied filtering (Supplementary Figure 1). Using the publicly-deposited dose-filtered tilt series from Figure 2, we generated two volumes: one reconstructed with the weighted-back projection method implemented in AreTomo3 [45] without additional processing, and another with an applied Wiener-like deconvolution filter for contrast enhancement [41] (Supplementary Figure 1A,D). Both volumes were segmented using MemBrain-seg [39] (Supplementary Figure 1B,E) and analyzed with the thickness measurement workflow (Supplementary Figure 1C,F). The analysis revealed only minor variations in the absolute thickness values between the two inputs (typically 0.2-0.3 nm). Importantly, the relative thickness patterns observed in the segmentation from the denoised tomogram were consistent. In both cases, the IMM was overall thicker compared to the OMM, and the outer chloroplast membrane was thicker than the thylakoids. This consistency demonstrates the robustness of the computational workflow, while underscoring the importance of consistent input data processing for reliable quantitative comparisons of membrane thicknesses.

### Consistent organelle-specific membrane thickness differences across *Chlamydomonas reinhardtii* cells

To investigate whether the organelle-specific membrane thickness patterns observed in a single tomogram (Figure 2) represent consistent, potentially biologically-relevant features, we extended the analysis to 51 denoised tomograms from the EMPIAR-11830 *Chlamydomonas reinhardtii* dataset [37]. We analyzed tomograms from different acquisition sessions for each membrane type of interest: nuclear envelope (NE, 10 tomograms), endoplasmic reticulum (ER, 18 tomograms), outer and inner mitochondrial membranes (OMM and IMM, 19 tomograms each), thylakoid (14 tomograms), and chloroplast membranes (14 tomograms for the thylakoid, 9 tomograms for the outer envelope membrane). Using instance segmentations with unique membrane labels as inputs, we generated millions of point-to-point distance (thickness) measurements per membrane type. To compare thickness consistency across tomograms, we plotted the median thickness values of each membrane type within each analyzed tomogram (Figure 3). The Golgi apparatus represents a special case, where we plotted three median thickness points per tomogram to capture the systematic thickness variations across different cisternae positions within the same organelle (examined in more detail below and in Figure 5).

**Figure 5.**
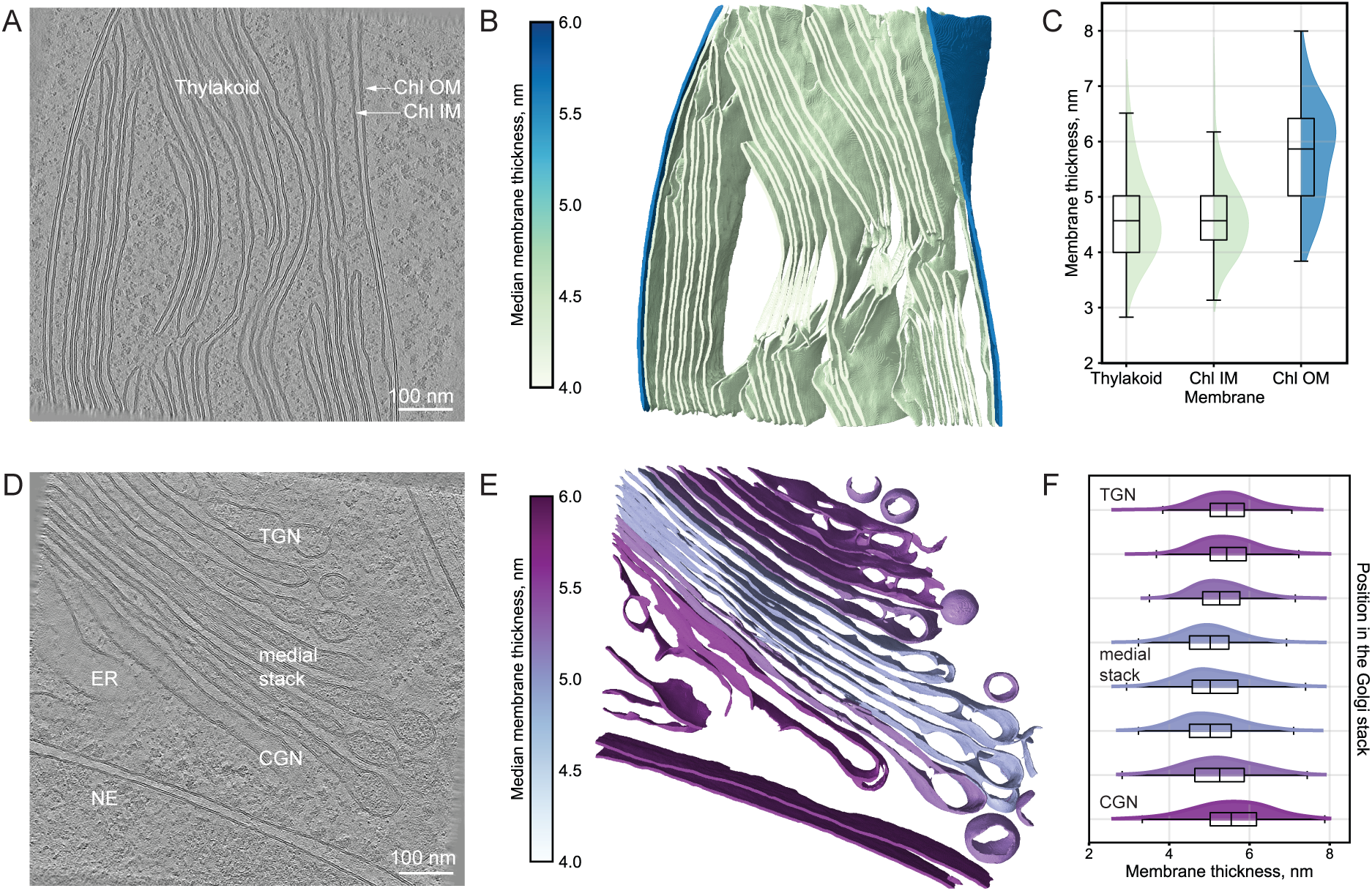
Membrane thickness variations within compartments of the same organelle. **A-C** Analysis of the chloroplast membrane system. **A** Central slice from *Chlamydomonas reinhardtii* tomogram [37] (EMPIAR-11830, 573.mrc, 7.84 Å/pixel at bin4), showing the thylakoid, inner and outer chloroplast membranes (Chl IM and Chl OM). **B** Thickness-mapped segmentation of the chloroplast, where color values indicate median membrane thickness. **C** Comparative thickness distributions show that the outer chloroplast membrane has significantly greater thickness compared to both the inner chloroplast membrane and the thylakoid. **D-F** Analysis of the Golgi apparatus membranes. **D** Central slice from *Chlamydomonas reinhardtii* tomogram [37] (EMPIAR-11830, 1055.mrc, 7.84 Å/pixel at bin4), showing the nuclear envelope (NE), endoplasmic reticulum (ER), and Golgi cisternae from the cis Golgi network (CGN) through the medial Golgi stack to trans Golgi network (TGN). Scale bar, 100 nm. **E** Thickness-mapped segmentation of the membranes, where color values indicate median membrane thickness. **F** Thickness distribution analysis across the Golgi stack highlights distinct cisternae populations within the Golgi apparatus. The membrane thickness progresses from ER-like values in the CGN, to a decrease in thickness in the medial cisternae, followed by a thickness increase in the TGN. Coated vesicles are similarly thick to the ER and CGN/TGN rather than the medial stack.

We found that NE and ER membranes exhibit the highest median thickness values (5.3-6.2 nm), followed by the outer chloroplast membrane (5.0-6.2 nm), the Golgi (4.7-5.9 nm), the thylakoid (4.5-5.0 nm), and mitochondrial membranes (OMM: 4.2-4.8 nm, IMM: 4.6-5.1 nm).

Statistical analysis revealed consistent patterns across tomograms: while NE and ER membranes had similar median thicknesses (unpaired t-test, not significant), paired t-tests on per-tomogram basis showed statistically-significant differences in the median thicknesses between the OMM and IMM (p < 0.001), the thylakoid and outer chloroplast membranes (p < 0.001), and the ER and medial Golgi membranes (p < 0.001, empty circles on Figure 3).

To further verify the reliability of the measurements, we examined whether the thickness distributions for a given membrane type remained consistent across tomograms where the membrane orientation and imaging conditions varied. Using the NE as a test case, we found that the median thickness values and the shapes of the underlying distributions remained nearly identical across tomograms (Supplementary Figure 2). The consistency of these results indicates that the analysis workflow is robust to variations in membrane geometry. This observation suggests that the thickness patterns in Figure 3 likely reflect genuine biologically-relevant properties of the studied membranes, as membrane thickness variations are more pronounced between different organelles than between individual cells.

### Human cells exhibit similar thickness relationships across membrane types

To further validate our workflow and to determine whether the organelle-specific thickness patterns observed for *Chlamydomonas reinhardtii* represent a more general principle of membrane biology, we acquired a smaller tomographic dataset of human embryonic kidney (HEK293) cells. These tomograms were processed using the described workflow. While there are small variations in the absolute thickness values between human and *Chlamydomonas* cells, we observe similar relative thickness relationships between different membrane types and organelles (Figure 4A). The NE and ER are consistently thicker than the mitochondrial membranes. Within mitochondria, the IMM is consistently and significantly thicker than the OMM (paired t-test, p = 0.002), in line with the relationship observed for *Chlamydomonas*.

These findings demonstrate two important points: first, the computational workflow can be applied to tomograms containing different membrane types across species; and second, relationships between the thicknesses of certain membranes appear to be conserved across human and algae cells. The general nature of these thickness patterns across evolutionarily distant species may suggest functional requirements that dictate similar distributions of lipids and membrane proteins across cellular compartments.

We hypothesized that the consistent membrane thickness relationships might in part reflect their different lipid compositions, as suggested by lipidomic analysis of subcellular fractions. To test whether our approach could detect lipid composition-dependent changes in membrane thickness, we performed a pharmacologic perturbation experiment using methyl-β-cyclodextrin (MBCD), a compound that extracts cholesterol from the plasma membranes (PM) of cells [46]. Compared to statins that lower cholesterol levels on long time scales [47], MBCD causes acute cholesterol depletion and is often used for cellular assays [46][48]. MBCD-treated cells were incubated with the compound dissolved in serum-free media for 30 minutes, while control cells were exposed to serum-free media alone for the same time prior to sample vitrification by plunge freezing. Tilt-series were acquired on thinned lamellae targeting the PM. Analysis of four PM instances per condition revealed a shift towards higher membrane thickness values in cholesterol-depleted cells (Figure 4B). The effectiveness of cholesterol depletion was assessed in parallel using a fluorometric detection assay on cells from the same passage, treated under identical conditions. This assay showed an approximate 40 % reduction in cholesterol levels, normalized to either soluble protein content or cell count (Supplementary Table 2). These findings demonstrate that our workflow can detect subtle changes in membrane thickness following lipid composition perturbation experiments.

### Membrane thickness variations within compartments of single organelles

The distinct functional roles of membranes within the same organelle are often reflected in their lipid and protein compositions, which likely account for the thickness variations we measured between the IMM and OMM in both *Chlamydomonas reinhardtii* and human kidney cell tomograms (Figure 4; see discussion). In *Chlamydomonas* chloroplasts, the outer envelope membrane (median thickness ≈ 5.7 nm) was consistently thicker than both the inner envelope and thylakoid membranes (median thicknesses ≈ 4.5 nm, Figure 5A-C). These variations align well with known differences in lipid composition between these membranes (see discussion). Additionally, local thickness maps revealed that closely spaced thylakoid membranes display regions of higher thickness and clustering of photosystem II complexes (Supplementary Figure 3A-C).

The Golgi apparatus represents another striking example for intrinsic variations within a single organelle. Composed of a series of flattened membrane-enclosed cisternae and vesicles, it functions as the primary sorting center for protein and lipid trafficking, directing cargo from the ER to the PM and endolysosomal system. Our analysis of the *Chlamydomonas* Golgi revealed a membrane thickness gradient across the stack (Figure 5D-F). The membranes of the cis-Golgi network exhibit low lumenal protein density, similar to ER membranes, and have comparable median thickness values (≈ 5.5 nm). Progressing through the flattened medial Golgi stack, lumenal protein density increases, and several consecutive cisternae membranes display a consistent reduced median thicknesses of ≈ 5 nm. This pattern aligns with observations that specific glycosylation enzymes distribute across two to three cisternae in specific sub-Golgi regions, depending on their position in the glycosylation pathway [49]. In the trans Golgi network, where protein sorting for PM delivery occurs, median membrane thickness increases to ≈ 5.4 nm, coupled with a decrease in lumenal protein density. The membrane thickness values of coated vesicles resemble those of the ER and trans Golgi cisternae, rather than the medial stack. This quantitative thickness analysis provides independent structural evidence supporting functional specialization across different regions of the Golgi apparatus, in line with the membrane thickness sorting hypothesis, proposed by Munro and colleagues [25][26][29][49] (see discussion).

## Discussion

In this study, we present a semi-automated computationally efficient workflow for measuring local membrane thickness from cryo-electron tomogram segmentations, implemented in publicly available Python code that can be integrated into existing cryo-ET analysis pipelines. Our analysis of *Chlamydomonas reinhardtii* and human cell tomograms reveals systematic thickness relationships across organelle membranes. Specifically, we observe that the endoplasmic reticulum (ER) and nuclear envelope (NE) show similar thickness profiles, consistent with their continuous nature and similar lipid compositions [22]. In contrast, mitochondrial and thylakoid membranes are consistently thinner than the ER, Golgi, and NE, possibly due to their specialized compositions and unique functional roles (Figure 3). These findings align with experimental reports on organelle-specific lipid compositions and provide a first quantitative analysis of membrane thickness heterogeneity in cellular contexts.

Our measurements are broadly consistent with previously reported bilayer thicknesses of compositionally less complex in vitro systems. Depending on the acyl chain length, the hydrophobic core thickness has been shown to vary between ≈ 2.5 and 4 nm, as measured by SAXS [33] and cryo-EM [34][35][36], while SANS measurements have reported thickness values for the total bilayer, including the water hydration layer, between ≈ 4.7 to 6.4 nm [31][32]. The cellular membrane thicknesses we obtain fall within this latter range, suggesting that the input membrane segmentations capture the full bilayer, including both the hydrophobic core and hydrophilic head groups. This is likely because U-Net-based segmentation models identify membranes based on contrast differences in electron scattering potential, particularly between the electron-dense, negatively charged phosphate groups in the head regions, the surrounding aqueous environment on one side, and the uncharged hydrophobic core on the other. Contrast in cryo-EM arises from interactions between the electron beam and the sample’s electrostatic potential [38], which is unevenly distributed across the lipid bilayer. This results in a distinct electron density profile [50], which segmentation models may learn to recognize in the reconstructed tomograms. If the hydration layer thickness (≈ 1.5 nm, as determined by SANS [31]) is added to SAXS- and cryo-EM-derived measurements of the hydrophobic core, the resulting values closely match the thickness distributions we observe. This agreement with established experimental methods suggests that our approach can be used to confidently measure membrane thickness in native cellular environments.

The workflow we propose differs from previous approaches in several key aspects: Instead of confining the analysis to 2D projections, we measure membrane thickness in 3D by computing Euclidean distances between paired surface points, guided by the direction of their normal vectors, thus accounting for local variations in membrane curvature. Furthermore, the thickness measurement step is parallelized over GPU nodes or CPU threads, enabling high-throughput analysis of hundreds of thousands to millions of point-to-point distance calculations per membrane within a tomogram. This high sampling density enhances the statistics of the resulting measurements and ensures their sensitivity to both global thickness differences across organelles and local variations within individual membranes. The consistent thickness distributions observed for a given organelle across tomograms (collected in different sessions under varying acquisition conditions, Supplementary Figure 2) demonstrate the method’s robustness. Finally, the resulting membrane thickness map can be directly overlaid with the input tomogram or with coordinates of complexes identified by subtomogram averaging or template matching, enabling contextual analysis of local membrane thickness variations in relation to protein localization.

Beyond organelle-wide differences, we detect systematic thickness variations within the membranes of individual organelles. The inner mitochondrial membrane (IMM) is consistently thicker than the outer mitochondrial membrane (OMM) within the same tomogram (Figure 2D, Figure 3). The properties of the IMM are thought to largely derive from its high cardiolipin content (≈ 20% of the lipid mass) – a diphosphatidylglycerol lipid with a small hydrophilic head group and large hydrophobic tail region that facilitate the high membrane curvature in cristae [23]. Additionally, the IMM is highly rich in protein, with respiratory complex proteins constituting ≈ 60-70 % of its mass [51]. Similar membrane specialization exists in chloroplasts, which are photosynthetic organelles enveloped by two membranes, whose distinct functions are dictated by the differences in lipid and protein compositions. Within the stromal volume, the protein complexes of photosystems I and II required for light absorption and ATP synthesis, sit on the thylakoid membranes, whose lipid composition is similar to that of the inner envelope membrane [52]. We observe that the outer envelope membrane is overall thicker than both the inner envelope and thylakoid membranes (Figure 5A-C). The inner envelope and thylakoid membranes exhibit overlapping thickness distributions, consistent with lipidomic studies that find that these two membranes have similar lipid profiles, while differing compositionally from the outer envelope membrane [52].

Perhaps most striking are the thickness gradients we observe across the Golgi apparatus. According to the membrane thickness sorting hypothesis proposed by Munro and colleagues, membrane protein localization throughout the secretory pathway primarily depends on the match between the length of the transmembrane span with the thickness of the bilayer [25][26][29][49]. This model posits that lipid composition and membrane thickness change accordingly: cis-Golgi cisternae should be surrounded by thin, phospholipid-rich membranes with low cholesterol content, similar to the ER, while trans-Golgi cisternae membranes should become progressively thicker as sphingolipid and sterol content increases [53]. Sphingolipid synthesis in the Golgi drives cholesterol enrichment, with cholesterol transported from the ER via non-vesicular mechanisms [53]. This increasing membrane thickness from the cis to trans Golgi would favor trafficking of proteins with longer transmembrane spans to the plasma membrane (PM), as supported by immunofluorescent experiments showing that PM proteins with synthetic transmembrane domains of 23 leucines reach the cell surface, while those with truncated domains (17 leucines) accumulate in the Golgi [29].

Our data provide structural evidence for key aspects of the membrane thickness sorting mechanism. We observe that membrane thickness increases in discrete steps from the medial Golgi towards the trans face: the medial cisternae exhibit higher lumenal density (likely representing Golgi-resident proteins) and thinner membranes compared to the trans cisternae, which have lower lumenal protein density and thicker membranes (Figure 5D-F). In agreement with the model, we also observe that the ER and cis Golgi cisternae display similar median thicknesses, although both are thicker than the medial Golgi. Despite its low reported cholesterol content, we consistently find that the ER is among the thicker membranes (second to the PM) in both *Chlamydomonas* and humans kidney cells (Figure 3 and Figure 4A). Adjacent to the ER, we observe that a few cis Golgi cisternae display intermediate median thicknesses that gradually decline towards the medial Golgi (Figure 5D-F). This pattern suggests that while the thickness-based sorting principle holds true across the Golgi stack, particularly onwards from the medial cisternae, additional factors might influence membrane content and thickness in the early secretory pathway. How these thickness relationships relate to the functional mechanism of protein sorting in the Golgi remains to be investigated.

Despite these promising results, our method has the following limitations: First, the input tomograms require sufficient signal-to-noise ratio for optimal segmentation results, and manual curation of instance segmentation labels may occasionally be necessary. Second, thickness estimation becomes less reliable in regions with extreme curvature changes (e.g. on self-folding membranes) due to reduced surface point coverage and potential inaccuracies in normal vector orientations. However, the high number of measurements across each membrane typically mitigates these local inaccuracies, maintaining statistical robustness. Third, the approach is constrained by the voxel size of input tomograms; therefore, thickness variations should be interpreted across sufficiently large membrane patches rather than at individual point pairs to avoid misinterpretation of measurement noise. Fourth, membrane thickness serves only as an indirect proxy for compositional differences across organelle membranes. Not only lipids, but also membrane proteins contribute electron density at the bilayer boundary, potentially influencing segmentation outcomes and, consequently, the thickness measurements. Indeed, our measurements likely reflect the combined contributions of lipids and integral proteins, both of which shape the biophysical properties of cellular membranes. A more comprehensive understanding of membrane organization could be achieved by using our method as a complementary approach to other techniques, such as high-resolution correlative light and electron microscopy workflows with chemically modified lipid probes [54].

At a finer scale, the lipid bilayer composition is thought to vary across the leaflets of a single membrane, a phenomenon known as lipid asymmetry, and laterally within membranes, referred to as microdomains [55]. While the ER maintains relative compositional equilibrium between its leaflets through ATP-independent transporters, the Golgi, PM, and endosomal membranes exhibit compositional asymmetry. In principle, our workflow could be expanded to include a lipid asymmetry and microdomain detection by systematically comparing the electron optical density between opposing leaflets.

Our cholesterol depletion experiment had been aimed at detecting effects on lipid asymmetry or microdomain organization. Instead, however, we observed an increase in PM thickness under cholesterol depletion conditions (Figure 4B). Although this result may be unexpected at first sight when compared to prior in vitro results [31], we note that cholesterol-depleted cells displayed morphological changes, likely indicating a complex multi-faceted cellular response to the acute treatment. Therefore, the observed thickening of the PM could reflect aberrant clustering of membrane proteins into patches on the cell surface due to significant perturbations of the bilayer, similarly to [56]. These findings do not rule out the presence of lipid asymmetry or microdomains, but suggest that further experiments would be required to better understand these phenomena in a complex cellular context.

Our workflow for measuring bilayer thickness from cryo-electron tomogram segmentations offers a systematic framework to quantify thickness variations of membranes in their native cellular contexts. It provides an orthogonal tool to address open questions in membrane biology. For example, building on our findings on the thickness gradients across the Golgi membranes, a similar approach can be applied to examine how disruptions in the protein sorting machinery affect membrane architecture in ER or Golgi trafficking mutants. Combined with perturbation experiments, our workflow can provide a direct readout of the effects of temperature changes, pharmacological treatments, or genetic modifications of lipid metabolism on membrane organization and thickness. Furthermore, integrating measurements of thickness variations within membrane microdomains with co-localization data for specific lipid and protein species, can offer an unbiased in situ approach to testing the lipid raft hypothesis. Finally, a community-driven effort could enable broader taxonomic sampling to assess whether the patterns observed in *Chlamydomonas reinhardtii* and human cells are conserved across different organisms. With its publicly available code and ease of implementation, we anticipate that our approach has the potential to become a valuable tool for the community, enabling researchers to probe questions related to membrane organization in situ.

## Acknowledgements

We thank the members of the Department of Molecular Sociology, in particular Beata Turoňová, Jenny Sachweh, Jan Philipp Kreysing, and Jiasui Liu for support and fruitful discussions on the project. M.B. acknowledges funding by the Max Plack Society.

## Author contributions

Conceptualization: D.G., M.B.; Methodology: D.G.; Formal analysis: D.G.; Visualization: D.G.; Writing – Original Draft: D.G., M.B.; Writing – Review & Editing: D.G., M.B., S.B.; Project Administration: M.B.; Supervision: M.B.; Funding Acquisition: M.B.

## Materials & Correspondence

Correspondence and material requests should be addressed to Martin Beck.

## Methods

### Cell culture

Human embryonic kidney HEK293 (HEK Flp-In T-Rex 293, Invitrogen) cells were cultured in DMEM (Sigma-Aldrich) supplemented with 10% fetal bovine serum (FBS, Gibco) under standard tissue culture conditions (37°C, 5% CO2) in T75 or T25 cell culture flasks (Greiner Bio-One).

### Pharmacological cholesterol depletion

Acute cholesterol depletion was induced in HEK293 cells using 10 mM methyl-β-cyclodextrin (MBCD, Sigma Aldrich). Cells were seeded on electron microscopy (EM) grids or in 6-well plates and incubated for 30 minutes in FBS-free DMEM supplemented with 25 mM HEPES buffer. Control cells were treated with FBS-free DMEM and 25 mM HEPES alone for the same duration. Following treatment, cells were either plunge-frozen for further cryo-EM sample preparation or washed and pelleted for cholesterol quantification.

### Cholesterol quantification

Cholesterol levels in MBCD-treated and control HEK293 cells were measured using a fluorometric assay kit (Amplex Red, Thermo Fisher Scientific) in duplicate for each condition. For this, HEK293 cells were seeded in 6-well plates overnight, treated as described, washed with PBS, trypsinized, and resuspended in DMEM. Cell suspensions were used to obtain cell counts using an automated counter (CellDrop, DeNovix). Cells were then gently pelleted in a pre-cooled centrifuge at 4°C and the cell culture media was removed. Cell pellets were lysed in detergent-containing buffer (kit-supplied) by vortexing, two freeze-thaw cycles, and gentle sonication. Soluble protein concentration was measured using the bicinchoninic acid (BCA) assay (Pierce BCA Protein Assay Kit, Thermo Fisher Scientific). Fluorescence in the lysed samples was generated by initiating enzyme-coupled reactions following cholesterol oxidation (all following the manufacturer’s instructions) and measured in light-impermeable 96-well plates with an automated plate reader (Tecan). Cholesterol concentrations in the cell samples were determined by normalizing fluorescence values to a standard curve. Cholesterol levels were normalized to either soluble protein content or cell count (Supplementary Table 2).

### Cryo-electron microscopy sample preparation

EM support grids (Au, 200 mesh, R2/2, SiO_2_ foil, Quantifoil) were glow discharged using a Pelco easiGlow discharger (15mA, 90 s). The foil-side of the grids was functionalized by incubation with 30 µg/mL laminin (Roche) for 1 h at room temperature or overnight at 4°C. Excess laminin solution was aspirated, the grids were washed twice with PBS and placed in a 35 mm cell culture dish containing DMEM and FBS. Adherent HEK cells (3-5 × 10^4^ cells/mL) were seeded on the foil side of grids and allowed to attach overnight. Cells were treated as described, back-blotted to remove excess media, and vitrified by plunge freezing into liquid ethane with a Leica EM GP2 plunger. The grids were assembled into AutoGrid cartridges (Thermo Fisher Scientific). To obtain electron-transparent sections of the sample, lamellae were prepared by cryo-focused ion beam (FIB) milling on a dual-beam Aquilos FIB-scanning electron microscope (FIB-SEM) (Thermo Fisher Scientific), equipped with a gallium ion source, similarly to a previously-described protocol [57]. Briefly, samples were coated with an organometallic platinum layer via a gas injection system and sputter-coated with platinum (20 s, 1 kV, 10 mA). Automated rough milling was performed using the AutoTEM 5 Software (Thermo Fisher Scientific), with stepwise reductions in the milling current and the distance between milling patterns. Fine milling was performed manually at 10-30 pA with milling patterns spaced 130-200 nm apart. The milling process was guided by SEM imaging (13 pA 2- 10 kV). A final platinum sputter coat (2 s,1 kV, 10 mA) was applied at the end of the session.

### Cryo-electron tomogram acquisition

The tilt series used for membrane thickness analysis in control and cholesterol-depleted HEK293 cells were acquired from three grids across three acquisition sessions (two for control and one for cholesterol-depleted cells). Data was collected on a Titan Krios G4 transmission electron microscope, equipped with a cold field emission gun and Falcon 4i direct electron detector, operated at 300 kV acceleration voltage in counting mode (Thermo Fisher Scientific). Suitable acquisition areas on each lamella were selected with overview images with 3.0 nm pixel size. Tilt series were collected using a dose-symmetric scheme [58] in 2° increments, grouped by two, over a ±60° range with a target cumulative electron dose of 120–130 e⁻/Å². Projection images were recorded at 64k magnification, corresponding to pixel size of 1.97 Å, with 10-eV wide energy slit inserted and nominal defocus varied in 0.5 µm steps in the range of −2.5 to −4.5 µm. Each projection image was acquired in low dose mode as a 4096 × 4096 pixel 10-frame movie, with frames motion-corrected and aligned on the fly in SerialEM [59].

### Tomogram reconstruction

Tilt series were preprocessed using the cryoKIT_10_ toolkit that combines multiple common tools used for cryo-ET processing. Individual tilt images were visually assessed, and those with substantial ice reflections, drift, or lamella edge obstructions were excluded. Additionally, tilt images with defocus values (estimated with Gctf v1.06 [60]) that deviated significantly from the mean were discarded, regardless of perceived visual quality. The remaining tilt images were dose-filtered based on cumulative electron dose [61], as previously described [62]. Cleaned, dose-filtered tilt series were aligned using patch tracking in AreTomo2 (for human kidney cells) or AreTomo3 (for *Chlamydomonas* tomograms) [45]. Tomograms were reconstructed at bin4 using the weighted back-projection (WBP) method [63], implemented in AreTomo.

### Contrast enhancement via denoising and deconvolution filtering

To enhance contrast and improve segmentation accuracy of HEK293 cell tomograms, denoising was performed using the cryoCARE U-Net [40], applied to even and odd movie frames. Tomograms were then reconstructed using the WBP, as described. CryoCARE-denoised *Chlamydomonas reinhardtii* tomograms were directly obtained from EMPIAR-11830 [37]. To evaluate the impact of contrast enhancement on membrane segmentation accuracy (Supplementary Figure 1), a WBP-reconstructed *Chlamydomonas* tomogram was further processed using a Wiener-like deconvolution filter implemented in Warp [41].

### Instance membrane segmentation

Membrane segmentations were generated using MemBrain-seg U-Net with the pre-trained MemBrain_seg_v10_alpha.ckpt model, applying the connected components analysis option to generate unique labels for individual membrane instances within the tomographic volume [39]. In certain cases, segmentation labels were manually curated in napari [42] by using the seg- select plugin [64] to merge segmentation labels (e.g., to combine individual cristae in a single label for IMM). The instance segmentation labels were assigned to their respective membrane instances based on human judgment. For visualization segmentations were rendered in ChimeraX and color-coded according either to their membrane identity (Figure 1, Figure 2, Supplementary Figure 1) or to their median membrane thickness (Figure 5).

### Membrane thickness analysis

The procedure for membrane thickness measurements was described in detail in the main text in section “Outline of the computational workflow for in situ membrane thickness analysis” and is visualized on Figure 1. The complete <u>Python source code</u> (memthick.py), a <u>detailed</u> <u>tutorial</u>and <u>documentation</u>, are available in the <u>cryoCAT</u> GitHub public repository [65].

### Statistical analysis

Membrane thickness distributions on Figure 2E were analyzed using complementary statistical approaches. We applied the Kolmogorov-Smirnov (KS) test to assesses differences between the entire distribution shapes rather than just their central tendencies. The KS test produces a D-statistic representing the maximum distance between cumulative distribution functions, with corresponding p-values, interpreted using standard significance thresholds: (ns, p > 0.05, *: p < 0.05, **: p < 0.01, ***: p < 0.001). Given the large sample sizes contributing to each distribution, even small differences could achieve statistical significance. To measure distribution differences in biologically meaningful units, we calculated the Wasserstein distance, which represents the minimum "cost" of transforming one distribution into another. This metric is expressed in nanometers, providing an intuitive measure of distribution separation. We further quantified effect sizes using Cohen’s d, calculated as the difference between distribution means divided by the pooled standard deviation. Following standard conventions, effect sizes were interpreted as negligible (<0.2), small (0.2-0.5), medium (0.5- 0.8), or large (>0.8). To assess thickness differences between mitochondrial membranes and the ER, we concatenated distance measurements from the IMM and OMM in a combined list. All statistical analyses were performed using the scipy.stats package. Detailed results of the statistical analysis are provided in Supplementary Table 1.

For analyzing the median thickness values plotted on Figure 3, we employed paired or unpaired t-tests. For each tomogram, we first calculated the median thickness value for a given membrane label from thousands to millions of individual distance measurements, treating the tomogram as the experimental unit. For functionally-related membranes acquired within the same tomographic volume (OMM vs IMM, thylakoid vs chloroplast outer membrane, and ER vs medial Golgi), we used paired t-tests comparing these tomogram-level median values. For membrane types observed in different tomograms (NE vs ER), we applied unpaired t-tests using the same tomogram-level medians. The resulting p-values were interpreted using standard significance thresholds (ns: p > 0.05; *: p < 0.05; **: p < 0.01; ***: p < 0.001). The data points on the plot represent median thickness values for each membrane type per tomogram (NE, n=10; ER, n=18; OMM, n=19; IMM, n=19; thylakoid, n=14; Chl OM, n=9). For the Golgi apparatus (n=6 tomograms), median thicknesses were calculated for each individual cisterna.

## Data availability

The previously published tilt series and cryoCARE-denoised bin4 tomograms for *Chlamydomonas reihardtii*, <u>EMPIAR-11830</u>, are accessible through the Electron Microscopy Public Image Archive. The tilt series and cryoCARE-denoised bin4 tomograms for HEK293 cells used in this study will be made available through the Electron Microscopy Public Image Archive upon publication. The previously published structures for *Chlamydomonas* ATP synthase, EMD-51802, and photosystem II complex, EMD-51731, are available through the Electron Microscopy Data Bank and in the public repository: https://github.com/Chromatin-Structure-Rhythms-Lab/ChlamyAnnotations/tree/master/10.1101-2024.12.28.630444 (“densities” folder) [44]. The annotated particle positions and orientations for ATP synthase and photosystem II are available through the public repository: https://github.com/Chromatin-Structure-Rhythms-Lab/ChlamyAnnotations/tree/master/10.1101-2024.12.28.630444 (“star” folder) [44]. Source data will be provided with this paper upon publication.

## Code availability

The source code used in this study is part of the cryoCAT (Contextual Analysis Tool for cryo- electron tomography) public GitHub repository [65], specifically available at https://github.com/turonova/cryoCAT/blob/main/cryocat/memthick.py. A Jupyter notebook, outlining the usage of the code along with detailed documentation, can be found at https://cryocat.readthedocs.io/latest/tutorials/membrane_thickness/measure_thickness_tutoria l.html and https://cryocat.readthedocs.io/latest/generated/cryocat.memthick.html.

