## Supplementary Files for "In situ evidence for systematic membrane thickness variation across cellular organelles"

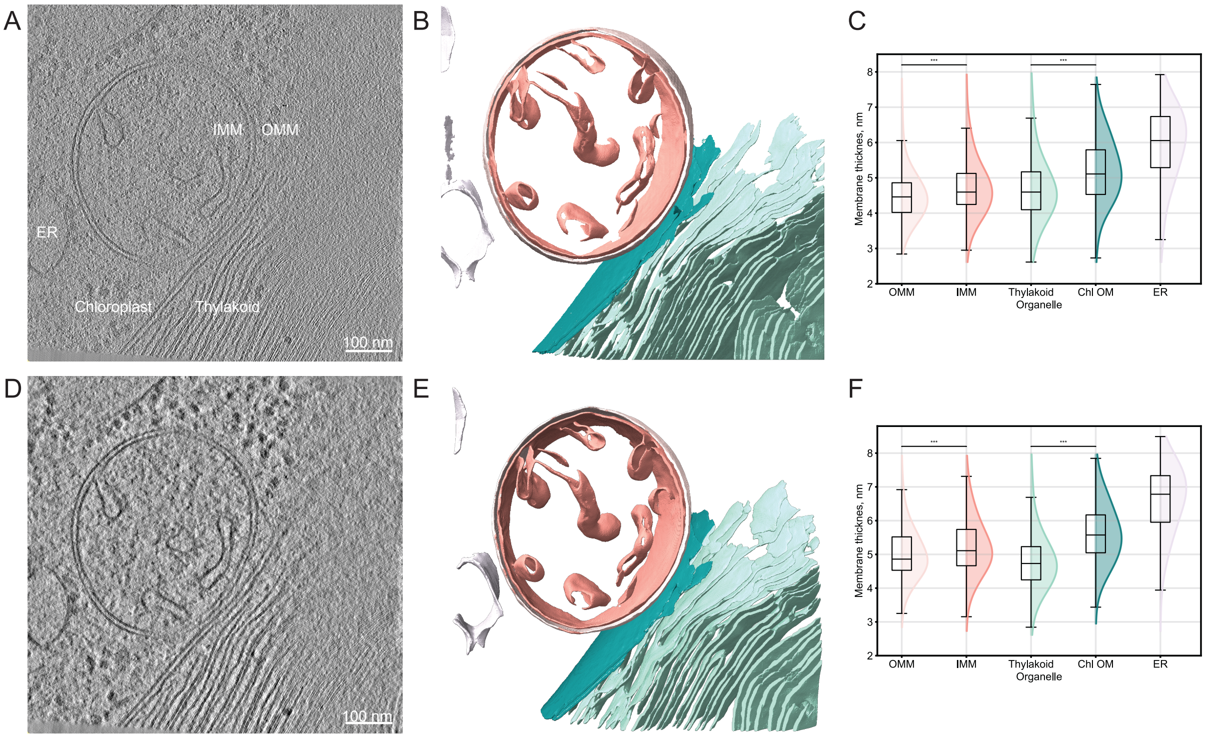


**Supplementary Figure 1 Impact of tomogram reconstruction and preprocessing on membrane thickness measurements. A-C** Analysis using tomogram reconstructed with weighted-back projection (WBP) method in AreTomo3 [45][63]. **D-E** Analysis of the WBP-reconstructed tomogram tomogram with applied Wiener-like deconvolution filter (Warp implementation [41]). **A,D** Central slices from both tomograms (7.84 Å/pix at bin4), showing the endoplasmic reticulum (ER), inner and outer mitochondrial membranes (IMM and OMM), thylakoid and chloroplast membranes. **B,E** MemBrain-seg [39] instance segmentation with distinct colors marking different membranes (color-coded as in Figure 2B). **C,F** The distribution plots show consistent thickness patterns for both input tomograms with statistically significant differences between OMM vs IMM and thylakoid vs chloroplast outer membrane (Chl OM) for both inputs, as determined by the Kolmogorov-Smirnov test (***: $p < 0.001$). While absolute median thickness values vary slightly between inputs (e.g., OMM ≈ 4.5 nm in WBP vs ≈ 4.8 nm with deconvolution), the relative relationships between membranes remain consistent. Scale bars, 100 nm.

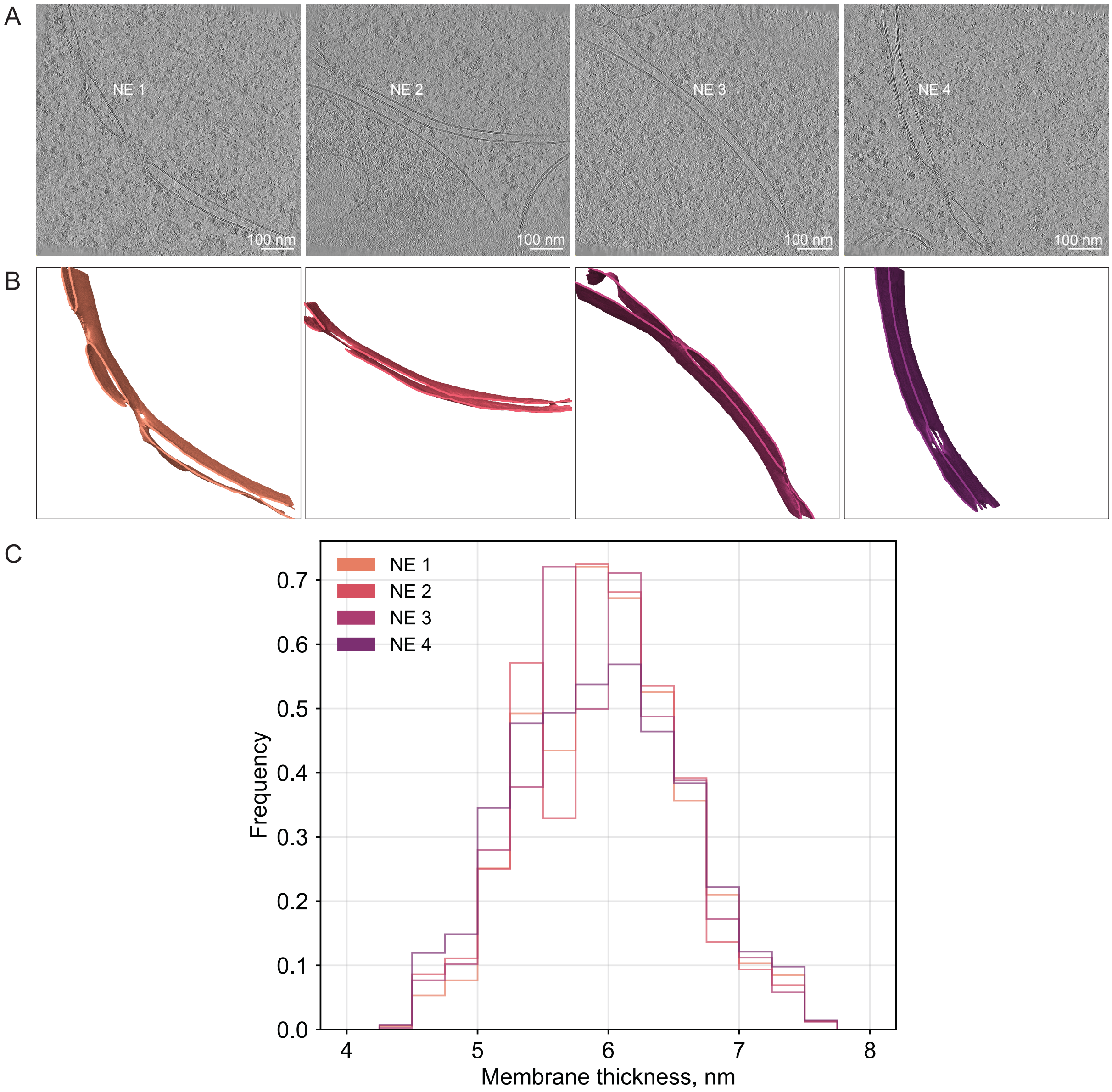


**Supplementary Figure 2 Consistent membrane thickness measurements across tomograms and membrane orientations, demonstrated for the nuclear envelope. A** Central slices from four *Chlamydomonas reinhardtii* tomograms showing nuclear envelopes in different orientations [37] (EMPIAR-11830, 7.84 Å/pixel at bin4, NE 1: 426.mrc, NE 2: 495.mrc, NE 3: 144.mrc, NE 4: 2174.mrc). Scale bars, 100 nm. **B** MemBrain [39] segmentations of the corresponding nuclear envelopes, colored to match distribution plots in C. **C** Thickness distribution histograms for the four nuclear envelopes – the measurements are highly consistent despite the different membrane orientations and the variation in imaging conditions.

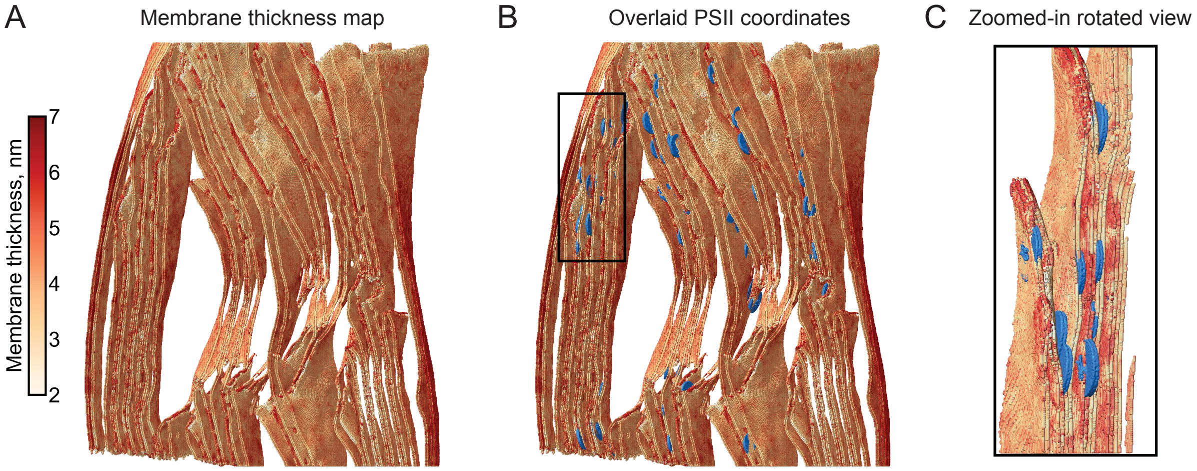


**Supplementary Figure 3 Local membrane thickness of the photosynthetic membranes from Figure 5A-C overlaid with the coordinates of photosystem II (PSII) complexes**. **A** Membrane thickness map for the thylakoid, chloroplast outer and inner membranes, with color intensity representing local membrane thickness. **B** Same view with overlaid PSII complex coordinates (blue) obtained from a publicly-available repository [44] as determined by subtomogram-averaging in [37], with the resulting subtomogram average (EMD-51731) used for visualization. **C** Zoomed-in rotated view of the region marked by black box in B, showing closely-spaced thylakoid membranes with increased thickness and clustered PSII complexes.

**Supplementary Table 1 Detailed statistical comparisons of the organelle membrane thickness distributions on Figure 2E.** For each membrane type, the sample sizes (total number of point-to-point distance measurements), as well as median and mean ± standard deviation (SD) thickness values are reported. The Wasserstein distance was used to quantify the average distance (in nm) required to transform one thickness distribution into another with 95% confidence intervals (CI) determined by bootstrap resampling. The Kolmogorov-Smirnov (KS) test was used to compare the thickness distributions with corresponding p-values indicating statistical significance. The magnitude of difference between distributions was quantified by Cohen's d, with effect sizes interpreted as negligible (<0.2), small (0.2-0.5), medium (0.5-0.8), or large (>0.8). Abbreviations: OMM (outer mitochondrial membrane), IMM (inner mitochondrial membrane), Chl OM (chloroplast outer membrane), ER (endoplasmic reticulum). For the comparison between mitochondrial membranes and ER, measurements for IMM and OMM were concatenated in a single list.

| Membrane instance | Per individual membrane instance | | | Paired statistical comparisons | | | |
|---|---|---|---|---|---|---|---|
| | Sample size | Median thickness, nm | Mean thickness ± SD, nm | Wasserstein distance, nm (95% CI) | p-value (KS test) | Effect size (95% CI) | Interpretation |
| OMM | 245 743 | 4.50 | 4.53 ± 0.67 | 0.2 (0.18, 0.23) | < 0.001 | -0.29 (-032, -0.27) | Small effect |
| IMM | 346 000 | 4.64 | 4.72 ± 0.72 | | | | |
| Thylakoid | 1 489 217 | 4.57 | 4.70 ± 0.81 | 0.62 (0.57, 0.64) | < 0.001 | -0.75 (-0.78, -0.72) | Medium effect |
| Chl OM | 82 030 | 5.26 | 5.32 ± 0.87 | | | | |
| ER | 85 189 | 5.71 | 5.64 ± 0.90 | 1.00 (0.97, 1.01) | < 0.001 | -1.22 (-1.25, -1.18) | Large effect |
| OMM+IMM | 591 743 | 4.57 | 4.64 ± 0.71 | | | | |

**Supplementary Table 2 Validation for cholesterol depletion in HEK293 cells via a fluorometric assay for the membrane thickness analysis shown on Figure 4B.** HEK293 cells were incubated for 30 minutes either in FBS-free DMEM containing 25 mM HEPES buffer (control, 0 mM MBCD) or with the same medium containing 10 mM MBCD to deplete cholesterol. Cholesterol levels were determined via a fluorometric assay (AmplexRed, Thermo Fisher Scientific) according to the manufacturer's protocol, with measurements performed in duplicate for each condition. Soluble protein concentration was determined by BCA assay. The table shows total cell counts, soluble protein concentration, and cholesterol levels in each condition, along with normalized values (cholesterol per protein [µg/µg] and µg cholesterol per cell). When normalized to protein content or per cell, treatment with 10 mM MBCD reduced cholesterol levels by 40%.

| | **0 mM MBCD** | **10 mM MBCD** | **[10 mM MBCD]/ [0mM MBCD]** |
|---|---|---|---|
| **Cells, N×$10^5$** | 3.4 | 4.1 | - |
| **Soluble protein, ×$10^2$ µg/mL** | 5.8 | 6.6 | - |
| **Cholesterol, µg/mL** | 8.7 | 6.1 | - |
| **[Cholesterol, µg] / [Soluble protein, µg], ×$10^{-2}$** | **1.5** | **0.9** | **0.6** |
| **[Cholesterol, µg] / [N cells], ×$10^{-6}$** | **2.1** | **1.2** | **0.6** |

**Supplementary Video 1:** Detailed visualization of the membrane thickness map shown in Figure 2C, along with overlaid coordinates of ATP synthase (blue), presented in Figure 2F. The video includes zoomed-in views and rotations of specific regions of the model. Membrane thickness is color-coded according to the scale in Figure 2C, with lighter colors indicating thinner membrane regions and darker reds representing thicker areas.
